# Unraveling the Genetic Comorbidity Landscape of Alzheimer’s Disease

**DOI:** 10.1101/2024.03.05.583453

**Authors:** Xueli Zhang, Dantong Li, Siting Ye, Shunming Liu, Shuo Ma, Min Li, Qiliang Peng, Lianting Hu, Xianwen Shang, Mingguang He, Lei Zhang

## Abstract

Alzheimer’s disease (AD) has emerged as the most prevalent and complex neurodegenerative disorder among the elderly population. However, the genetic comorbidity etiology for AD remains poorly understood. In this study, we conducted pleiotropic analysis for 41 AD phenotypic comorbidities, identifying ten genetic comorbidities with 16 pleiotropy genes associated with AD. Through biological functional and network analysis, we elucidated the molecular and functional landscape of AD genetic comorbidities. Furthermore, leveraging the pleiotropic genes and reported biomarkers for AD genetic comorbidities, we identified 50 potential biomarkers for AD diagnosis. Our findings deepen the understanding of the occurrence of AD genetic comorbidities and provide new insights for the search for AD diagnostic markers.

**Highlights:** The present study has focused on the comorbidities associated with Alzheimer’s disease (AD) by constructing a landscape of these comorbidities at various levels, including diseases, genetics, and pathways.

1. The study findings reveal novel and significant pathways that contribute to the etiology of AD and its comorbidities.
2. By exploring pleiotropic genes and reported biomarkers of AD comorbidities, the study has identified several potential diagnostic biomarker candidates for AD.

Graphic abstract.
Study pipeline.

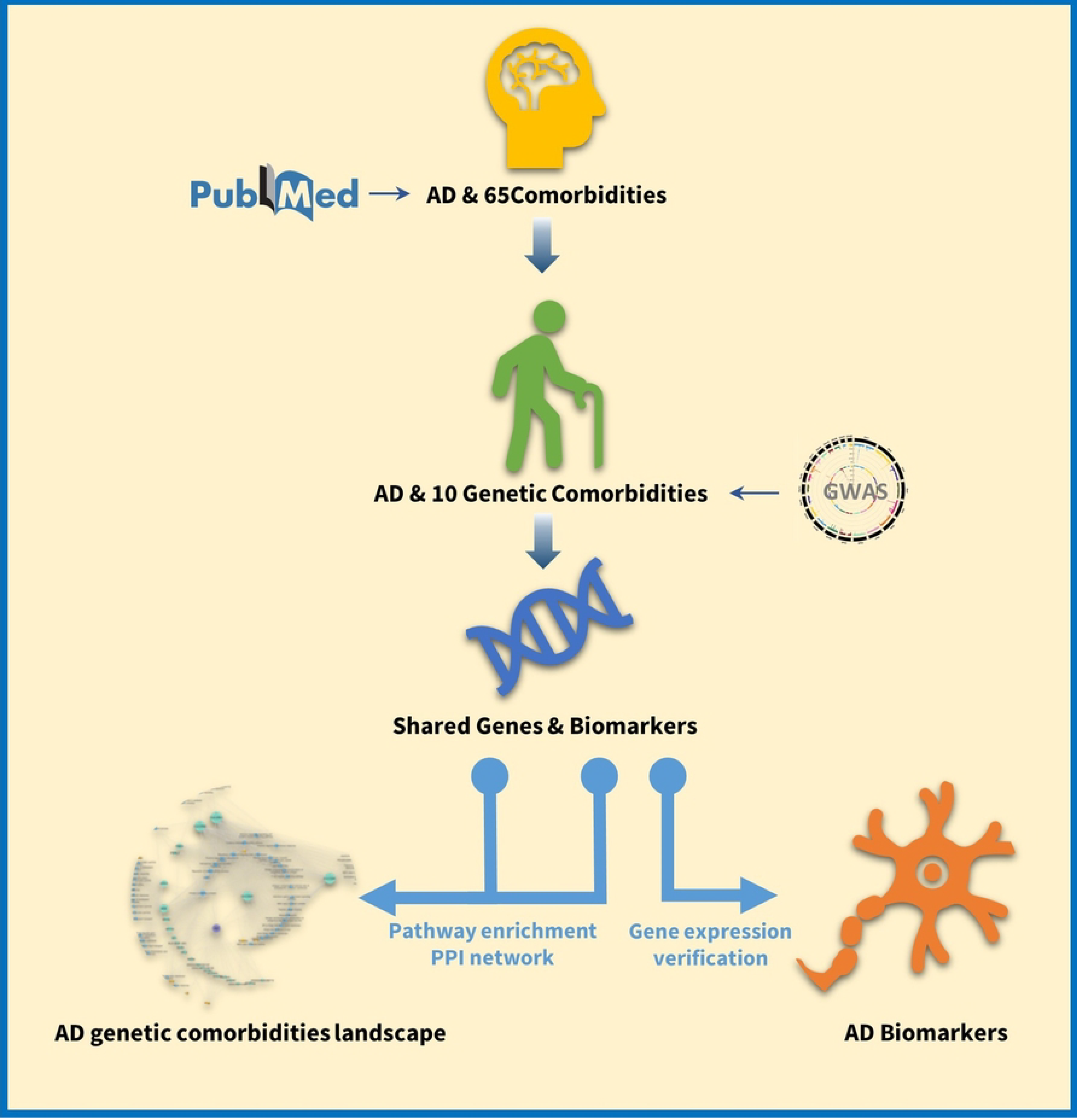

## Introduction

Alzheimer’s disease (AD) has emerged as the foremost prevalent and intricate neurodegenerative disorder among the geriatric populace(1, 2). Owing to the current lack of effective ways to intervene AD progression, it is critical to identify key determinants for the etiology and early diagnosis of AD.

Against the backdrop of a rapidly aging global population, the co-occurrence of comorbidities among individuals diagnosed with AD has emerged as a prominent and pervasive phenomenon(3). Considering the pervasive prevalence of AD and its profound implications for affected individuals, there exists an escalating imperative to meticulously investigate the intricate interplay between specific comorbidity patterns and their intricate association with AD. It is well-established that many chronic diseases precede the onset of AD, and their presence significantly elevates the risk of AD development. Consequently, exploring the etiology of comorbidities in relation to AD holds promise for advancing preventive measures and early diagnosis of AD. Moreover, the biomarkers identified for these comorbidities hold great potential as biomarkers for AD itself. As such, greater attention has been focused on investigating the association between comorbidity and AD(4-7).

Although several longitudinal studies have identified chronic diseases as at-risk conditions for increased AD incidence, most research on the association between AD and comorbidities has focused on the impact of a single or a small number of chronic diseases. This approach overlooks other frequent co-occurrences, thus restricting researchers from deeper exploration(4, 8, 9). Meanwhile, the majority of studies conducted thus far have primarily focused on exploring the phenotypic relationships between AD and its associated comorbidities. With the advent of Genome-wide Association Studies (GWAS), some genetic factors for the comorbidities associated with AD have been identified. For example, variants in genes involved in lipid metabolism, such as APOE gene and CLU gene, have been associated with an increased risk of AD and cardiovascular disease (2), and variants in the APOE gene, as well as genes involved in insulin signalling and glucose metabolism, have been implicated in the development of AD and type 2 diabetes(10). Nevertheless, the existing body of research has predominantly concentrated on investigating individual comorbidities, thereby leaving the comprehensive landscape of genetic comorbidities in AD largely unexplored.

Constructing the landscape of genetic comorbidities for AD could suggest a potential differential impact of specific comorbidity patterns on AD development and further provide more diagnostic biomarker candidates.

## Results

### Identification of AD genetic comorbidities

In accordance with the prescribed methods, a comprehensive literature search was performed resulting in the identification of 65 phenotypic comorbidities associated with AD (Figure 1). Of these, 44 displayed GWAS data, meeting the inclusion criteria for further examination. (Figure 1, Table S1) A total of 15,567,451 patients were included in this analysis. A meticulous quality control assessment was executed, and ultimately, a total of 39,391,465 SNPs were deemed suitable for further analysis. We used conditional QQ plots to detect the pleiotropy between AD and the comorbidities. Significant leftward shifts under various P value cut-offs (0.1, 0.01, 0.001, and 0.0001) were observed in QQ plots for eight diseases, including Crohn’s disease (CD), irritable bowel syndrome (IBS), chronic obstructive pulmonary disease (COPD), multiple sclerosis (MS), chronic sinusitis (CS), pernicious anaemia, chronic kidney disease, eczema (Figure 2). The QQ plots for the not significant comorbidities were shown in Figure S1.

**Figure 1.**
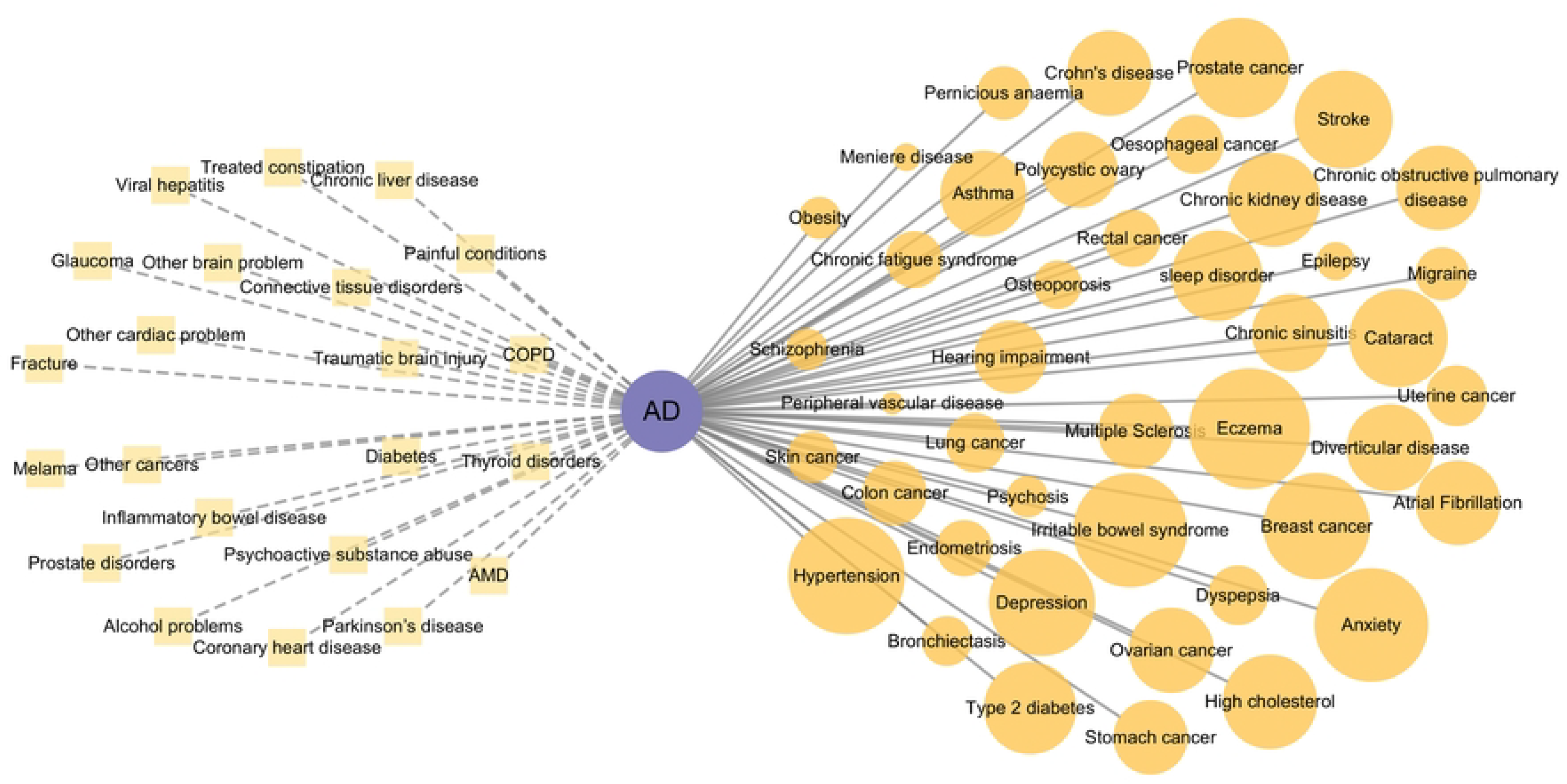
Knowledge graph for AD and its phenotypic comorbidities. 65 diseases were identified as phenotypic comorbidities for AD. The nodes represent the diseases, and the size of nodes indicate the number of samples. 44 diseases had GWAS data (circles), and the others not (blocks).

**Figure 2.**
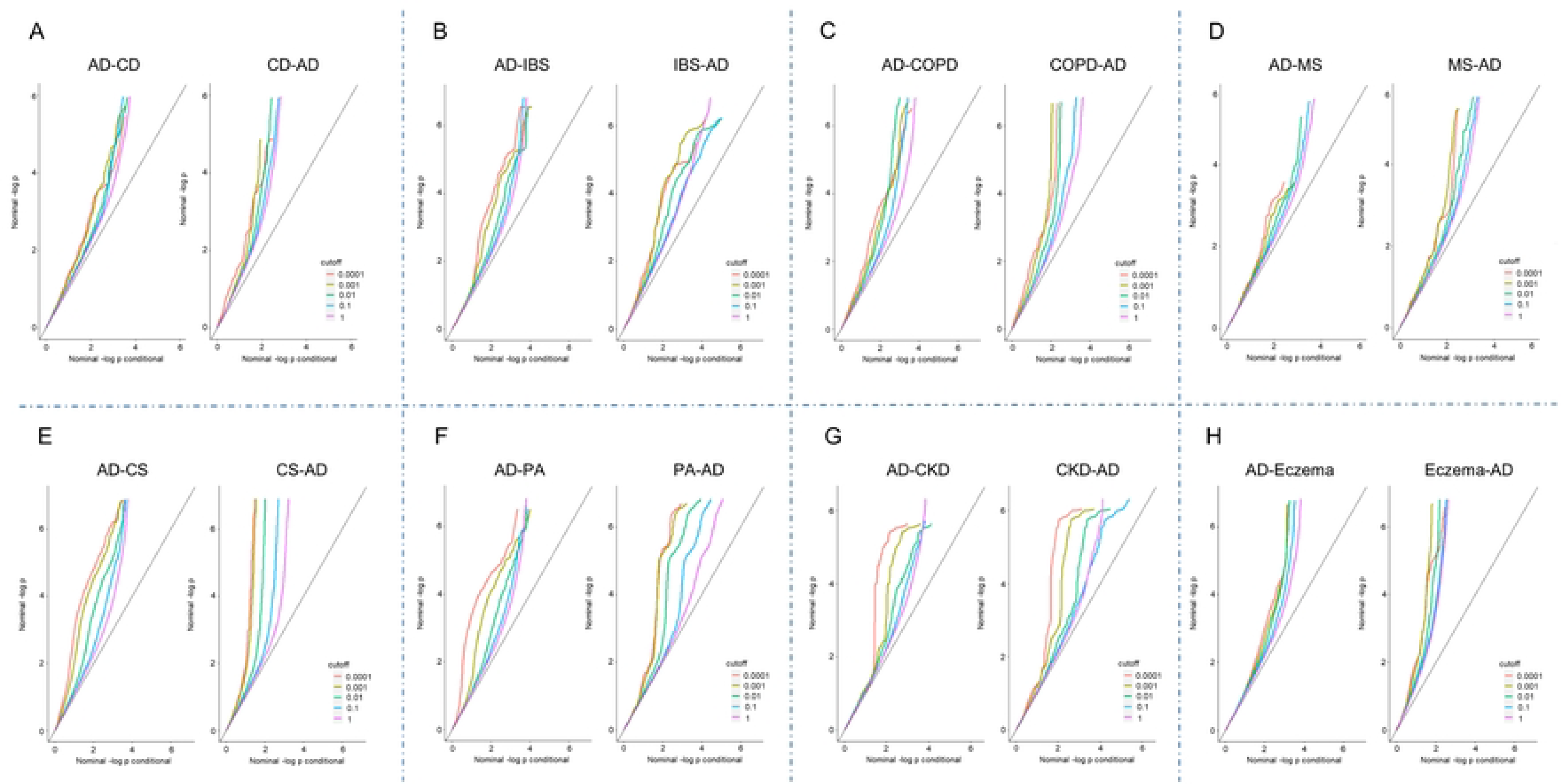
Conditional QQ plots for AD and its genetic comorbidities.

### Mapping of pleiotropic genes of AD-related genetic comorbidities

We plotted the genetic Manhant plots for AD and its genetic comorbidities in Figure 3, where the SNP situation for AD was presented in the inside circle and the comorbidities were in the outside circle. We found that chromosomal 6, 11, and 19 consist of the most significant shared loci for AD-related comorbidities. With predefined cut-off criteria (p<10^−5^ and ccFDR<0.01), we obtained 104 pleiotropic SNPs, which were mapped on 24 genes (Table S2). The pleiotropic SNPs for AD and CKD are all on Chromosome 19 and were mapped to MADD, ENSG00000255197, NR1H3, PSMC3, RAPSN and SPI1 gene. The pleiotropic SNPs between AD and COPD or CS or PA were all located on Chromosome 6 and mapped to HLA-DRB1, HLA-DQA1, HLA-DQB1, HLA-DRB5, HLA-DQB1-AS1 genes, respectively. All pleiotropic SNPs for AD and MS and some for AD and Eczema were on Chromosome 19 and mapped to CEACAM16-AS1, ENSG00000288773, NECTIN2, and BCL3 genes. SNPs for AD and Eczema were on Chromosome 6 and mapped to HLA-DRB1, and HLA-DQA1.

**Figure 3.**
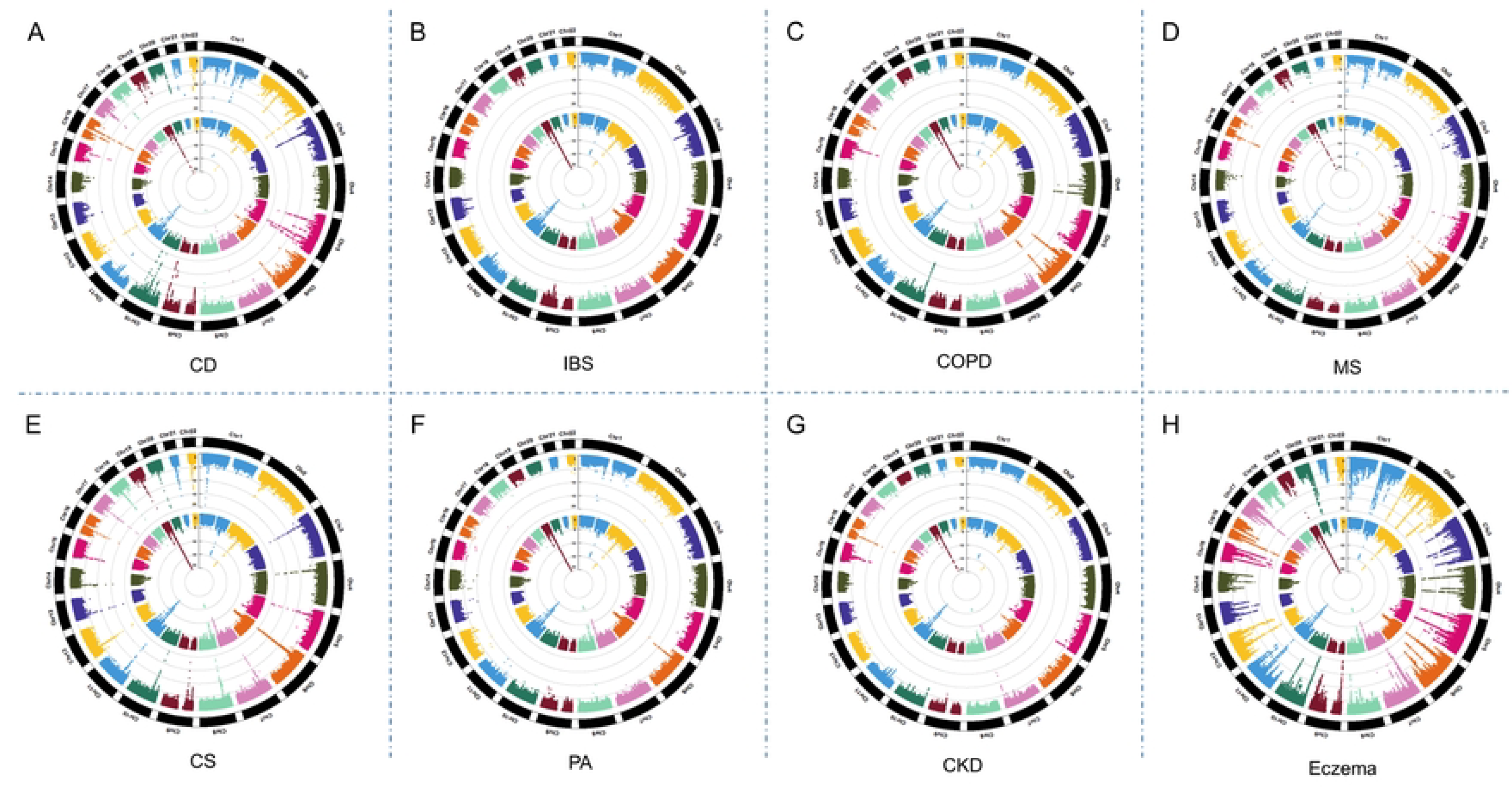
Manhattan plots for AD genetic comorbidities.

### Landscape of AD-related genetic comorbidities

Adding two previously reported genetic comorbidities, posttraumatic stress syndrome (PTSD) and age-related macular degeneration (AMD), we finally identified 10 genetic comorbidities for AD. We mapped the AD genetic comorbidities along with their pleiotropic genes, and pathways on one combined network, as the landscape of AD genetic comorbidities (Figure 4). The pleiotropic genes of AD-related comorbidities were significantly enriched in GO terms related to biological processes of immune response (e.g., very-low-density lipoprotein particle clearance, chylomicron remnant clearance, positive regulation of cholesterol esterification, MHC class II receptor activity) (Table S3). KEGG enrichment analysis demonstrated that the pleiotropic genes were mainly enriched in asthma, which is strongly mediated by pathways related to immunity. Genes in the HLA family, with the highest degree of interaction, were identified as hub genes (Figure 4, Figure S2).

**Figure 4.**
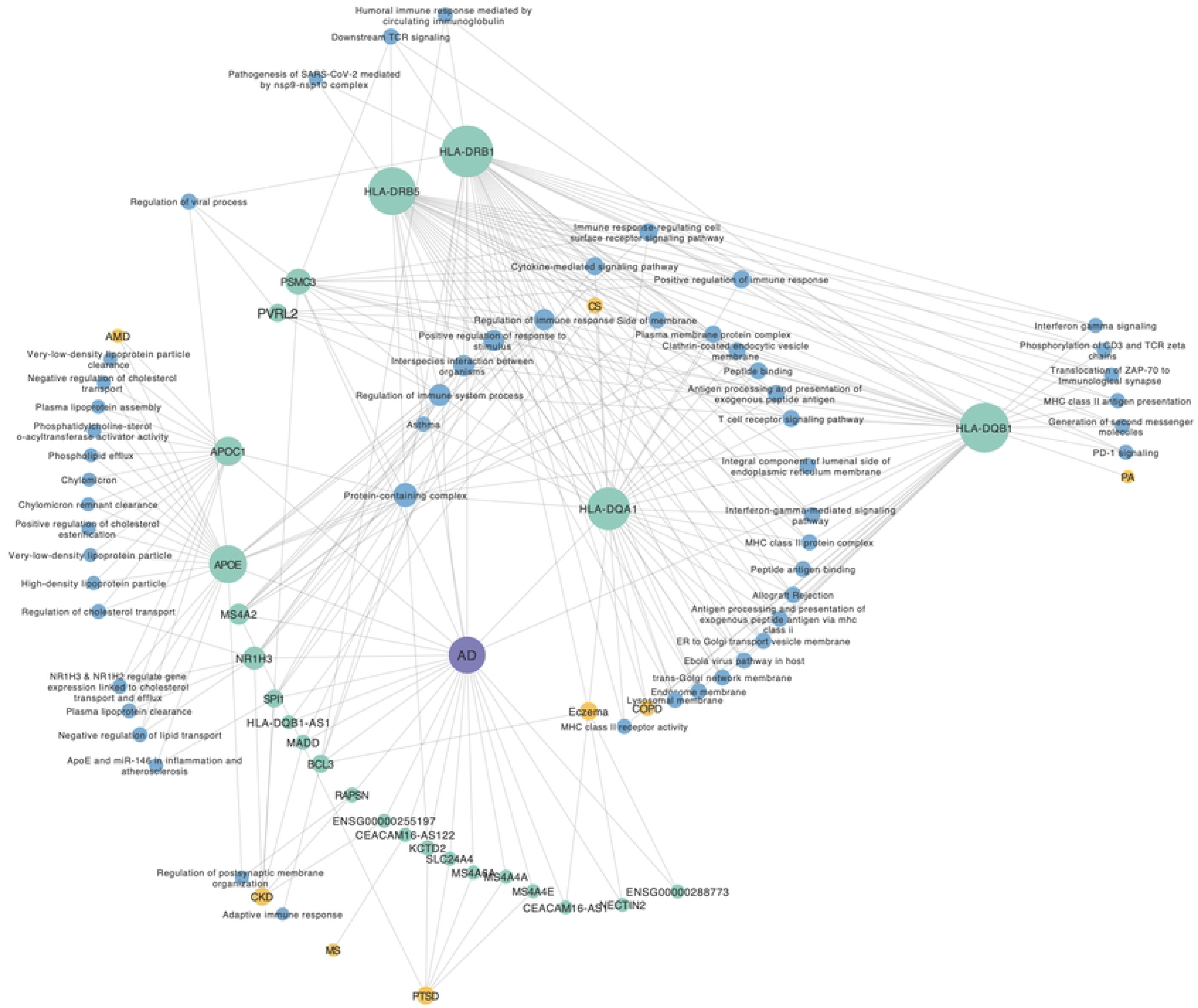
The landscape of AD and its genetic comorbidities. AD was set as a purple node. The yellow nodes represent genetic comorbidities for AD. The green nodes represent pleiotropic genes, and the blue nodes are enriched pathways by pleiotropic genes. The size represents the Degree of the nodes.

Based on further observation, COPD, CS, Eczema as well as PA tend to share genes, such as HLA-DQB1 and HLA-DRB1. In contrast, multimorbid relationships of CKD and AD, PTSD and AD tend to share immune-related pathways. The detrimental role of CKD on the brain has been previously reported, being CKD as a pro-inflammatory dysmetabolic state that is associated with brain dysfunction. SPI1, for example, a transcriptional activator that may be specifically involved in the differentiation or activation of macrophages or B-cells. MS4A2, the pleiotropic gene for PTSD and AD, was shown to mediate the secretion of important lymphokines. Although mediated by different genes, the identified comorbidities were all closely related to immune responses. Further, we also mapped the pleiotropic genes to the human PPI network and found that most of them connected closely with others (Figure 5A), indicating that they may conduct biological functions synergistically.

**Figure 5.**
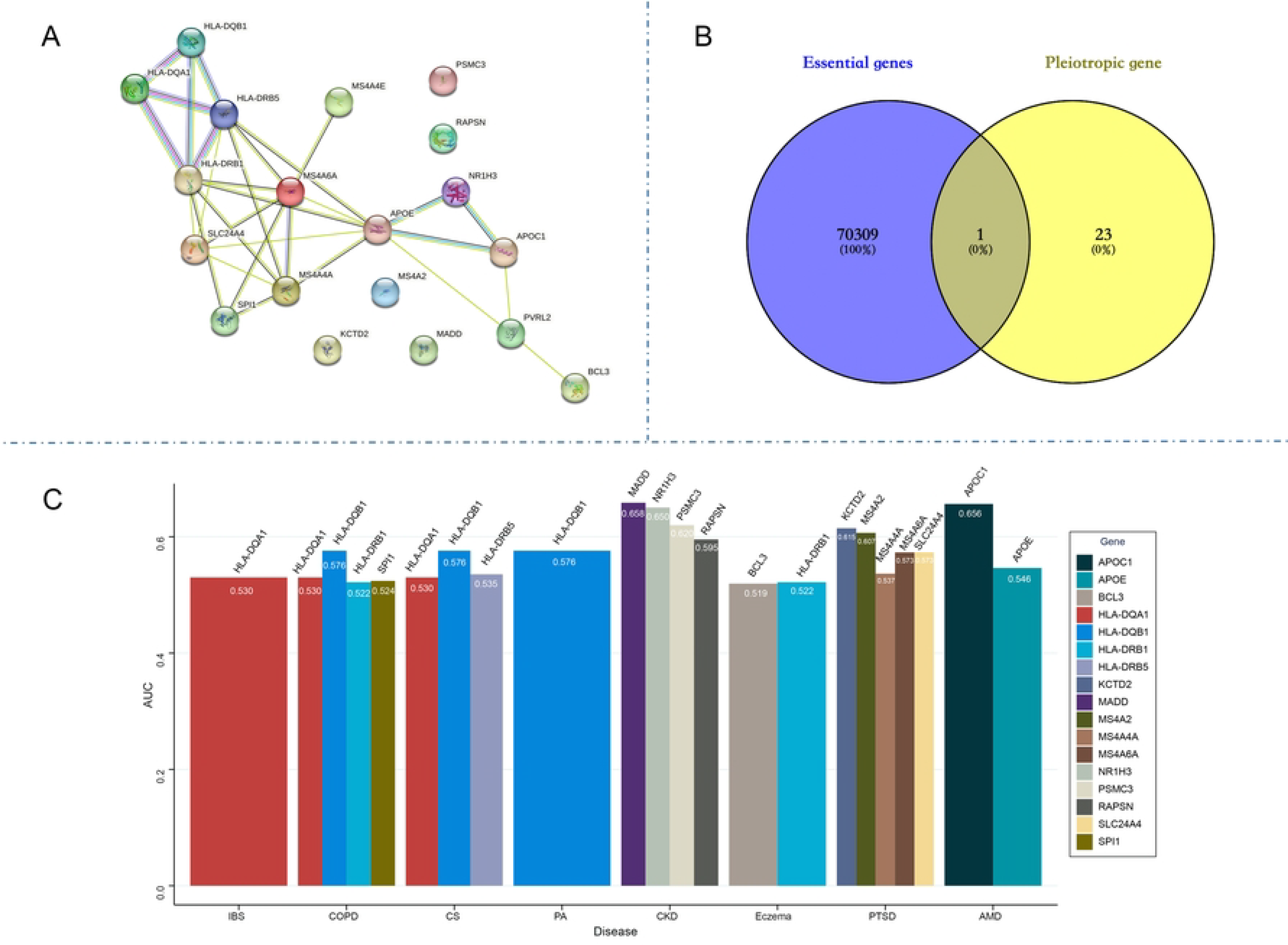
System analysis for pleiotropic genes. A. PPI network for pleiotropic genes. B. Venny plot for essential genes and AD pleiotropic genes. Only the SPI1 gene was both an essential gene and a pleiotropic gene. C. Diagnostic AUCs for pleiotropic genes. The x-axis shows the source comorbidities for the pleiotropic genes.

### Function and expression analysis for pleiotropic genes

Essential genes were reported to have a tendency to encode hub proteins in the human interactome and play important roles in maintaining normal developmental and/or physiological functions. It is curious that if pleiotropic genes could be essential genes. Here, we obtained 70310 essential genes, which were human orthologs of mouse genes whose disruptions are embryonically or postnatally lethal. We found that SPI1 was the only essential gene among pleiotropic genes (Figure 5B). These results indicated that most pleiotropic genes were functionally peripheral in the human interactome, and their mutations are compatible with survival into reproductive years so that these comorbidity phenotypes are preserved in a population. Housekeeping genes, also known as constitutive genes, are a class of genes that are expressed at relatively constant levels in all cells and under normal physiological conditions. These genes are responsible for carrying out fundamental cellular functions that are essential for the maintenance of basic cellular processes. To examine whether pleiotropic genes tend to be housekeeping genes, we summarized the number of tissues each gene was expressed in based on the gene expression data of 53 tissues in ScRNA-seq data from GTEx. We found that pleiotropic genes, such as APOE and HLA-DRB1 as well as PWMC3, tend to be expressed in more tissues (Figure S3).

### Identification of potential diagnostic biomarkers for AD from AD-related comorbidities

In order to find new biomarkers for AD, we tested the predictive efficiency of the above pleiotropic genes on two independent AD microarray datasets, respectively. Diagnostic test results for pleiotropic genes of different comorbidities have been presented in Figure 5C. APOC1 (the pleiotropic gene for AD and AMD, average AUC= 0.65), MADD, NR1H3, PSMC3 (the pleiotropic gene for AD and CKD, all average AUC >0.6), and KCTD2, MS4A2 (the pleiotropic gene for AD and PTSD, all average AUC >0.6) showed good diagnostic value in AD microarray datasets. However, the AUC values of the rest genes were not significant.

The number of the pleiotropic genes with predicted potential in our study was relatively low, we, therefore, searched the reported biomarkers of the 10 genetic comorbidities from public databases and examined their prediction accuracy (Table S4). Based on a logistic regression model, 50 genes passed the AUC of 0.8 on both validation datasets (Figure 6A). Notably, ACTB and YWHAZ showed good prediction accuracy for five different comorbidities (ACTB for CKD, COPD, CD, MS, PTSD and YWHAZ for COPD, CS, CD, Eczema and MS) while MSC for four comorbidities (COPD, CD, Eczema, MS). Interestingly, >10 of these biomarkers were mapped on Synapse, Cell junction, and Vesicle pathways in GO annotation on the cellular component level (Figure 6B). No overlap was found among pathways enriched by pleiotropic genes and biomarker candidates.

**Figure 6.**
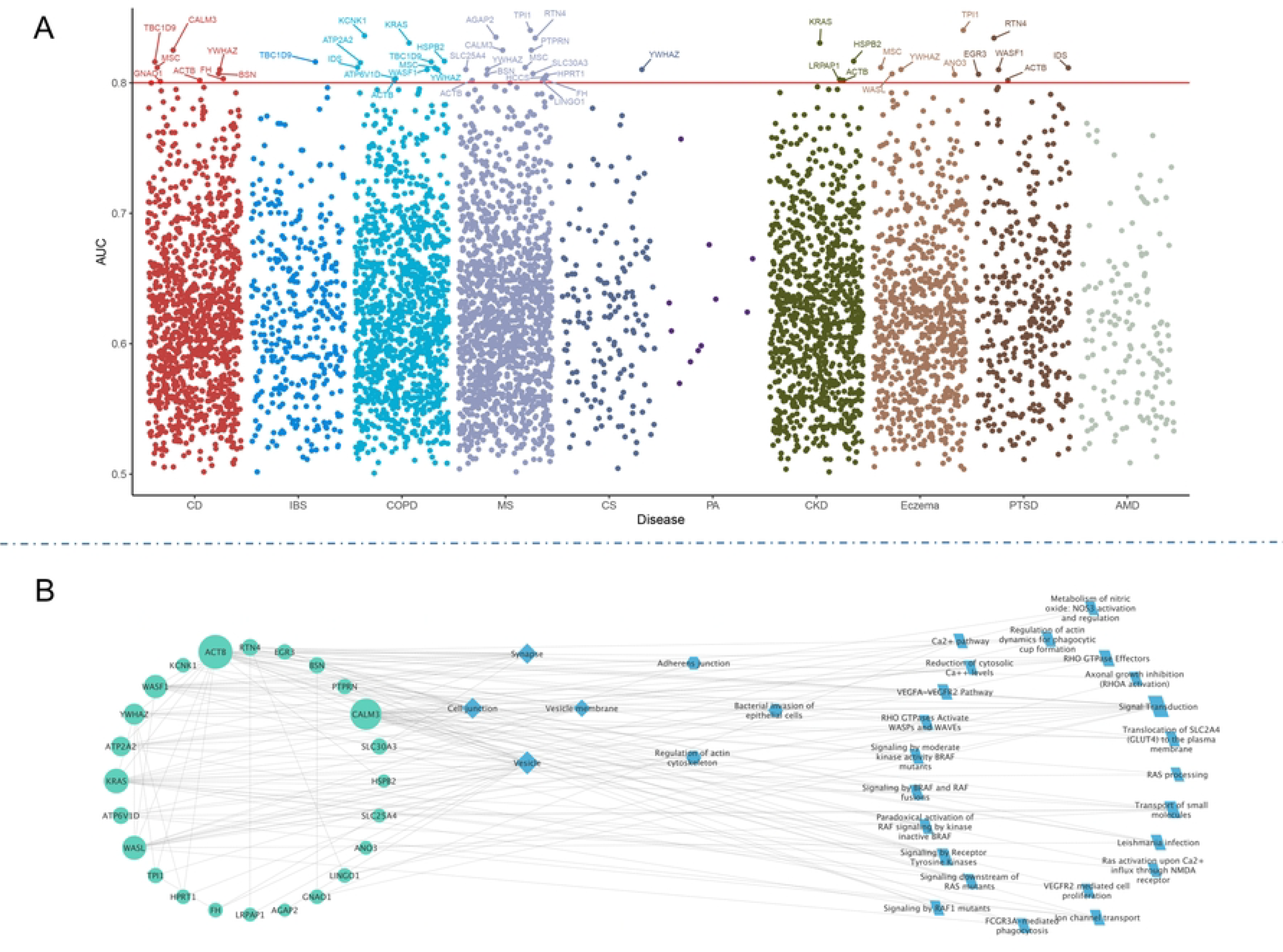
A. Diagnostic AUCs for genetics comorbidities biomarkers in AD. 50 biomarkers have an AUC>0.8, which were identified as biomarker candidates for AD. B. Combined network of PPI and pathway enrichment for biomarker candidates for AD. The circle nodes represent biomarker candidates, the diamond nodes represent GO pathways, the hexagon nodes are KEGG pathways, the parallelogram nodes are Reactome pathways, and the rectangle nodes represent Wiki pathways. The size of the nodes shows the Degree, the nodes with a bigger Degree have a bigger size.

## Discussion

Our study conducted a profiling of the genetic comorbidity relationships of AD, adding the functional and network analysis, we have formulated a comprehensive portrayal of the landscape of AD genetic comorbidities. Notably, this analysis represents the largest in scale to date. Further, we identified several novel diagnosis biomarkers for AD, from the pleiotropic genes and reported biomarkers of comorbidities. These findings portray a comprehensive genetic and multimorbid landscape for AD and reveal a pool of prospective biomarkers that may prove useful in the early diagnosis, management, and treatment of AD and its associated comorbidities.

Our study approved that AD was a hub comorbidity in old population. In light of the considerable prevalence and deleterious impact of AD, particularly among the elderly population, it is imperative that we direct our focus towards the comprehensive understanding and management of its comorbidities. The urgency to prioritize this matter stems from the recognition of the significant burden these comorbidities impose on overall health. Shang et al. found that individuals with ≥ six diseases were around four times more likely to develop dementia, and around 51.2% of incident dementia was attributed to one or more observed diseases(11). A few studies have investigated the association between comorbidity and incident dementia. However, these studies agree with our research highlighting the importance of comorbidity in the development of dementia. Shang et al. proved the association of high cholesterol with dementia(11). Giulia Grande et al. found increased dementia risk among persons with high levels of systemic inflammation(12). It is well acknowledged that a low-grade chronic proinflammatory state characterized by high levels of serum cytokines is common in older individuals and that older persons with high inflammatory markers have a higher number of chronic diseases as well as a steeper increase in comorbidity over time. The imbalance between inflammatory and anti-inflammatory agents due to several chronic diseases can lead to a systemic and chronic proinflammatory state, which ultimately may affect the brain.

In the present study, we constructed the landscape for AD and its comorbidity. Comorbidity has been found previously to be associated with biomarkers of neurodegeneration and amyloid deposition; however, specific comorbidity patterns may increase dementia risk through different pathways. The identified genetic comorbidity was attributed to genes encoding human leukocyte antigen (HLA) and major histocompatibility complex (MHC) class II receptor activity. HLA within the MHC in humans consists of several highly polymorphic and tightly linked genes on chromosome 6p21 (13). Multiple previous association studies verified that certain HLA gene variants within MHC class I and II regions have shown significant associations with AD, agreeing with our findings and indicating the shared patterns for the comorbidity (14, 15). A wide range of activities involved in immune responses may be determined by HLA genes, including inflammation, T-cell transendothelial migration, infection, brain development and plasticity in AD pathogenesis(13). HLA-DR, a known microglia marker for AD. The expression of HLA-DRB1 and HLA-DRB5 in the microglia has been proved positively correlated with measures of AD pathology(16). Furthermore, the immune response in the brain may be influenced by the peripheral immune system and vice versa, because the integrity of the blood–brain barrier (BBB) could be compromised by inflammatory processes and microvascular pathologies, both of which have been observed in AD(17, 18). It has also been demonstrated that macrophage phagocytosis can be impaired and HLA-DR can be abnormally expressed by neutrophils and monocytes(19, 20). Our study identified that Cholesterol Metabolism Pathway played an important role in the pleiotropy among AD and other combidities, which coincided a lasted study by Holstege et al.(21).

The present study exhibits a few noteworthy limitations. Firstly, despite the sample size being relatively large, it is important to note that the number of cases for each disease or multimorbid disease pair was limited. Consequently, it is possible that certain multimorbid disease pairs that are overrepresented in the population may have been overlooked. For instance, the cardiovascular diseases failed to achieve statistical significance in this study. Secondly, it is plausible that variants with minimal effects may have been overlooked by the GWAS analyses. Thirdly, although some biomarkers were identified in our analysis that are supported by previous literature, experimental validation is necessary to affirm their clinical utility.

## Conclusions

In summary, we have performed, for the largest scale, a systematic analysis of multimorbid relations for AD as well as their shared genetic components based on the GWAS analysis. Our findings reveal a propensity for comorbidity in AD patients and offer insight into the genetic mechanisms underlying these associations. Furthermore, the pleiotropic genes identified in our analysis may serve as valuable biomarkers for both AD and its comorbidities, enhancing the ability of researchers and clinicians to manage these conditions in a more holistic manner.

## Methods

### Literature search for AD phenotypic comorbidities

In the previous study, we identified 53 diseases associated with AD in phenotype(22). Then, a comprehensive literature search was conducted using the PubMed database to search other comorbidities identified by other researchers. The search employed the following keywords: “multiple diseases,” “multiple conditions,” “multiple chronic diseases,” “multiple chronic conditions,” “comorbidity,” “comorbidit*,” or “co-morbidit*,” combined with “dementia” or “Alzheimer’s disease.” No language restrictions were imposed. In total, until December 2023, 2350 published papers were identified, from which 65 phenotypic comorbidities associated with Alzheimer’s disease were extracted.

### GWAS data

We obtained GWAS summary statistics for Alzheimer’s disease (AD) and its associated comorbidities from the GWAS Catalog database. Table S1 provides detailed information about the acquired datasets. The AD dataset used in this study was sourced from Lambert et al. and consisted of a cohort comprising 17008 AD patients and 37154 controls without cognitive impairment.

### Estimation of pleiotropy among AD and its comorbidities

To assess pleiotropy among AD and its associated phenotypic comorbidities, we employed conditional quantile-quantile (QQ) plots. Pronounced pleiotropy was indicated by a leftward lean and noticeable separation among various cut-off points.

### Detection of pleiotropic genes among AD and its genetic comorbidities

In this study, we employed the conditional false discovery rate (cFDR) algorithm to identify pleiotropic genes associated with AD and its genetic comorbidities. We applied a rigorous threshold of statistical significance, using a cut-off value of 0.01 as a stringent criterion for detecting significant pleiotropy.

### Biological network and function analysis for pleiotropic genes

To determine whether the AD pleiotropic genes were essential genes, we used the OGEE database(23). Additionally, we utilized the String database to obtain protein-protein interaction and pathway information for the biological network and functional analysis of pleiotropic genes.

### Diagnostic biomarker discovery for AD

We downloaded gene expression data from the GEO database to identify new diagnostic biomarkers for AD. The dataset used for AD discovery (GSE36980) consisted of gene expression data obtained from the frontal cortex, temporal cortex, and hippocampus of 26 individuals with Alzheimer’s disease (AD) and 62 healthy individuals serving as controls. For AD replication (GSE132903), gene expression data from the middle temporal gyrus were analyzed, involving 97 AD patients and 98 healthy controls. The candidate biomarkers were derived from two sources: pleiotropic genes and previously identified biomarkers associated with genetic comorbidities of AD.

## Supplementary Data

Table S1. Data summary. A total of 65 phenotypic comorbidities were identified for AD, out of which 44 had summary statistics GWAS data available.

Figure S1. QQ plots for not genetically significant comorbidities.

Table S2. Pleiotropic SNPs and genes for AD.

Table S3. Pathways for pleiotropic genes.

Figure S2. Network topology analysis results for the landscape.

Figure S3. Expression of pleiotropic genes on bulk (A) and single-cell (B) RNA-seq.

Table S4. Reported biomarkers for AD genetic comorbidities.

## Acknowledgments

The authors are grateful to our colleagues and collaborators.

## Author Contributions

XZ, XS, LZ and MH contributed to the conception and design of the study; DL, SY, SL and LH contributed to the acquisition and analysis of data; SM and ML contributed to drafting the text or preparing the figures.

## Data Availability

All the included data could be visited and downloaded from open databases provided in Methods.

